# Medial orbitofrontal cortex, dorsolateral prefrontal cortex, and hippocampus differentially represent the event saliency

**DOI:** 10.1101/285718

**Authors:** Anna Jafarpour, Sandon Griffin, Jack J. Lin, Robert T. Knight

## Abstract

Two primary functions attributed to the hippocampus and prefrontal cortex network are retaining the temporal and spatial associations of events and detecting deviant events. It is, however, unclear how these two functions converge onto one mechanism. Here, we tested whether increased activity with perceiving salient events is a deviant detection signal or contains information about the event associations by reflecting the magnitude of deviance (i.e., event saliency). We also tested how the deviant detection signal is affected by the degree of anticipation. We studied regional neural activity when people watched a movie that had varying saliency of a novel or an anticipated flow of salient events. Using intracranial electroencephalography from ten patients, we observed that high-frequency activity (50-150 Hz) in the hippocampus, dorsolateral prefrontal cortex (dorsolateral PFC), and medial orbitofrontal cortex (OFC) tracked event saliency. We also observed that medial OFC activity was stronger when the salient events were anticipated than when they were novel. These results suggest that dorsolateral PFC and medial OFC, as well as the hippocampus, signify the saliency magnitude of events, reflecting the hierarchical structure of event associations.

## Introduction

‘I was waiting at home for my friend. I made some tea, washed the cups, and poured hot water. Then I felt everything shaking. It was an earthquake. I put the cup down and waited to see if there was an aftershock. Just about then, my friend arrived.’ We experience the world as a sequence of event, but we remember them as segmented sequences. During encoding, perceiving unusual events separates the flow of events, i.e., an event segmentation process (Zacks and Swallow, 2007; Zacks et al., 2007) so that each segment contains the relationship of events that occurred in a similar circumstance (Zwaan and Radvansky, 1998). In this process, sequences of events are organized to create a hierarchical relationship (Kurby and Zacks, 2008; Zacks and Swallow, 2007; Zwaan and Radvansky, 1998), where a sequence of less notable events are embedded in an overriding structure of segmented sequences (Hanson and Hirst, 1989, 1989). In this example, making some tea was embedded in a friend’s visiting. We hypothesized that a basic model to support the hierarchical relationship of sequences relies on the magnitude of event deviancy. In principle, the hierarchical relationship can be reconstructed from event saliency where less salient events are more temporally associated with prior events than more salient new events (Figure 1C, also see Yeung et al., 1996).

**Figure 1.**
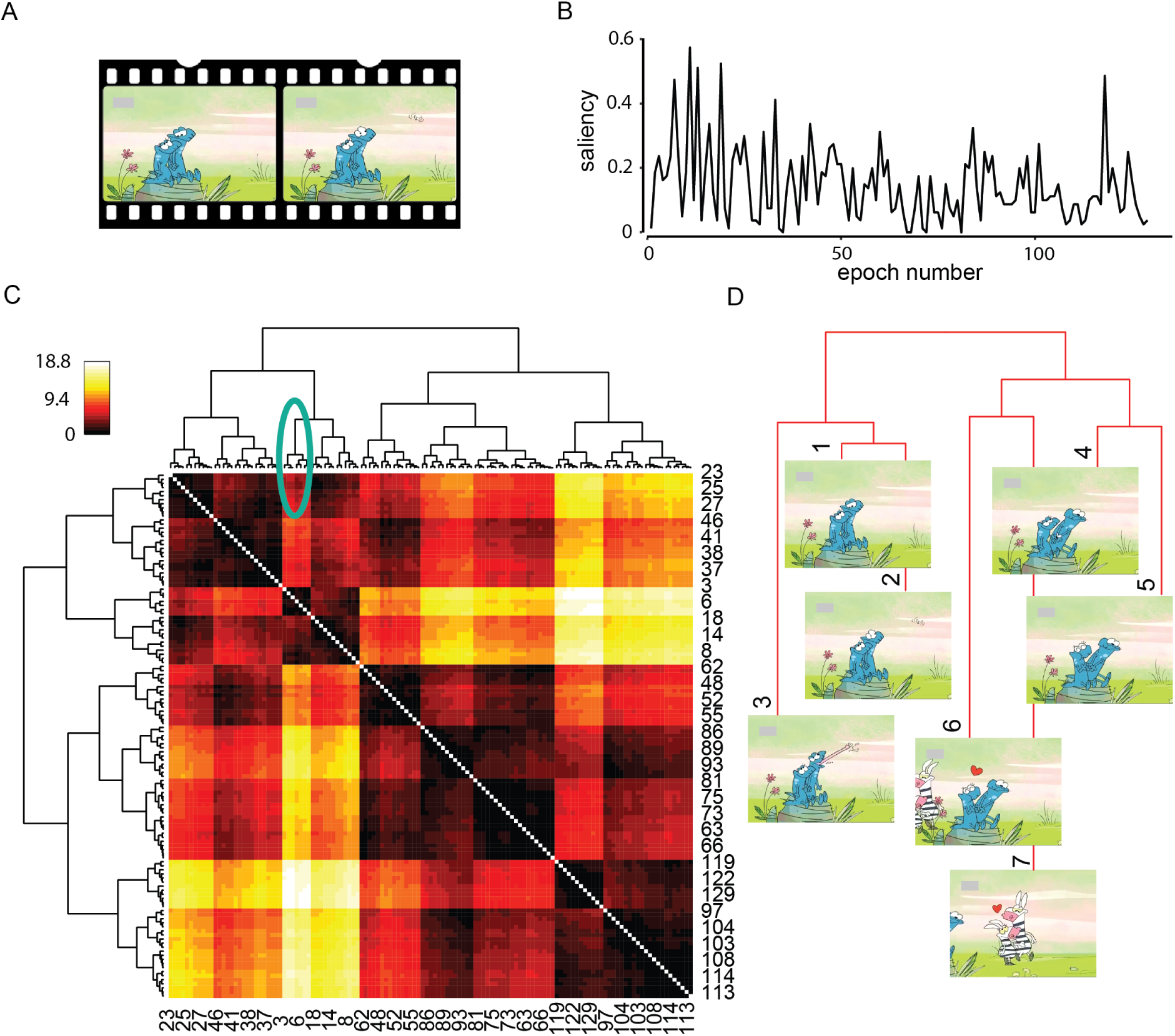
A hierarchical structure of the movie can be extracted from event saliency. (A) Patients passively watched a (∼ 3 minutes) muted animation that they did not see before. The movie had a mixture of novel and anticipated new events. (B) We chunked the movie into 129 interchangeable epochs of 1.5 seconds. The epochs had a range of event saliency as defined as the proportion of an independent group of participants (n = 80) that determined event-boundaries in the movie-epochs. (C) Event saliency was used to construct the hierarchical relationship of epochs, with distance being the sum of event saliency between epochs. The heat map shows the sum of saliency of the events that occurred between each pair of events (ranged between 0 and 18.8 that is the largest sum of saliency of a pair of events). The row and column order have been re-ordered based on the hierarchical clustering results. All the epochs are displayed in rows and columns (every third column is numbered in the illustration) and temporally adjacent events are next to each other. The right and left branches of the hierarchy do not imply any order and can be flipped since the plot is symmetric. (D) Zoomed in view of the first 10.5 seconds of the movie (marked branches in C). The graph shows the structure of event-associations at the start of the movie.

The magnitude of deviance of events is referred to as “event saliency” and is quantified by crowdsourcing. Previously event saliency has been used to determine the probability of a deviant event in a linguistic experiment (Coulson et al., 1998) but recently, with the availability of videos and advances in computer vision, this term is used for quantifying the magnitude of deviation in a flow of a movie. Event saliency is not determined by changes in a visual scene but relies on following the movies and noting significant changes in the flow of events (Rosani et al., 2015; Zhang et al., 2017). Accordingly, event saliency is measured by either asking an independent group to identify the boundaries or, in linguistics, studying the transition from one word to another, that is extracted from a large body of literature.

Both event association and deviancy detection are linked to the hippocampus, OFC, and dorsolateral prefrontal cortex (PFC) (Knight, 1996; Nobre et al., 1999; Paz et al., 2010; Zacks et al., 2007). We reasoned deviant detection reflects event saliency; this signal would also reflect a temporal association of events. Here we tested the prediction that high-frequency neural activity in subregions of the PFC and the hippocampus tracked event saliency.

The PFC-hippocampal neural network is also engaged in prospective coding (Brown et al., 2016; Hindy et al., 2016; Hsieh and Ranganath, 2015; Hsieh et al., 2014; Jafarpour et al., 2017) that enhances event segmentation (Schütz-Bosbach and Prinz, 2007). Note that anticipated salient events are different from novel events. For example, a salient event like a friend’s planned visiting is anticipated, whereas, a salient event such as an earthquake is novel. Here, we predicted that the neural representation of event saliency would be different for novel and predictable salient events.

Generating sequences of novel events in an experimental condition is challenging (Zarcone et al., 2016). Previous studies have used a discrete experimental design, comparing the neural activity at the time of perceiving deviant events, i.e., event-boundaries (Kurby and Zacks, 2008), to the neural activity at the time of perceiving non-boundary events (Speer et al., 2007; Whitney et al., 2009; Zacks et al., 2007). However, after encountering the first few deviant events participants anticipate the new events; thus, the later deviant events are no longer novel; instead, they become anticipated salient events. A flow of events, such as observed in a movie, has numerous event-boundaries with a range of anticipated or novel saliency providing an ideal experimental design to address this issue.

We recorded local field potential using intracranial electroencephalography (iEEG) from ten patients with epilepsy, who had electrodes implemented for clinical purposes. Patients passively watched a movie (Figure 1). The analysis focused on the local activity captured as the power in the high-frequency activity (HFA, 50 – 150 Hz) that serves as a metric for local neural activation (Belitski et al., 2008; Jacobs and Kahana, 2009; Lachaux et al., 2012; Ray et al., 2008a; Rich and Wallis, 2017).

## Materials and Methods

### Approval

The study protocol was approved by the Office for the Protection of Human Subjects of the University of California, Berkeley and the University of California, Irvine. All participants provided written informed consent before participating.

### Participants

#### Intracranial EEG

Ten epileptic patients who had stereo-tactically implanted depth electrodes to localize the seizure onset zone for subsequent surgical resection participated in this study, (4 female, mean age = 37, SD = 11, age range = 22 to 58 years old; Table 1). The electrodes were placed at the University of California, Irvine Medical Centre with 5-mm inter-electrode spacing. All patients had normal (or corrected) vision. No seizure occurred during task administration. Two independent neurologists inspected the neural activity and identified the electrodes with an epileptic activity which were excluded from the analysis so that all electrodes included in analysis were nonpathological and free of epileptogenic spikes. Any segment where focal spikes spread to other brain regions were also excluded from analysis (Table 1). Electrode coverage included the medial temporal lobe and the prefrontal cortex depending on their clinical requirements (Figure 2). Electrodes were localized in the patient’s native space, then transferred to MNI space for visualizing the group coverage. We studied electrodes in three regions of interests: the lateral PFC, the OFC, and the hippocampus (Table 1 and Figure 2).

**Table 1.** Patient electrode coverage. This table contains patients’ age (years old), gender (male or female), dominant hand (right or left), the number of bipolar-referenced electrodes in any region of the OFC, hippocampus, or lateral PFC, and the number of trials (out of 129 total trials).

| **Patient** | **Age** | **Gender** | **Dominant-hand** | **OFC** | **hippocampus** | **Lateral PFC** | **Number of trials** |
| --- | --- | --- | --- | --- | --- | --- | --- |
| **1** | 50 | Male | Right | 3 | 10 | 15 | 112 |
| **2** | 46 | Male | Right | 3 |  | 17 | 125 |
| **3** | 34 | Female | Right | 2 |  |  | 104 |
| **4** | 31 | Male | Left | 2 |  | 15 | 110 |
| **5** | 40 | Female | Left | 3 | 11 | 20 | 129 |
| **6** | 58 | Female | Right | 3 | 3 | 20 | 123 |
| **7** | 34 | Male | Right |  |  | 2 | 129 |
| **8** | 22 | Female | Right | 10 |  | 16 | 122 |
| **9** | 33 | Male | Right |  |  | 10 | 126 |
| **10** | 23 | Male | Right |  |  | 33 | 129 |

**Figure 2.**
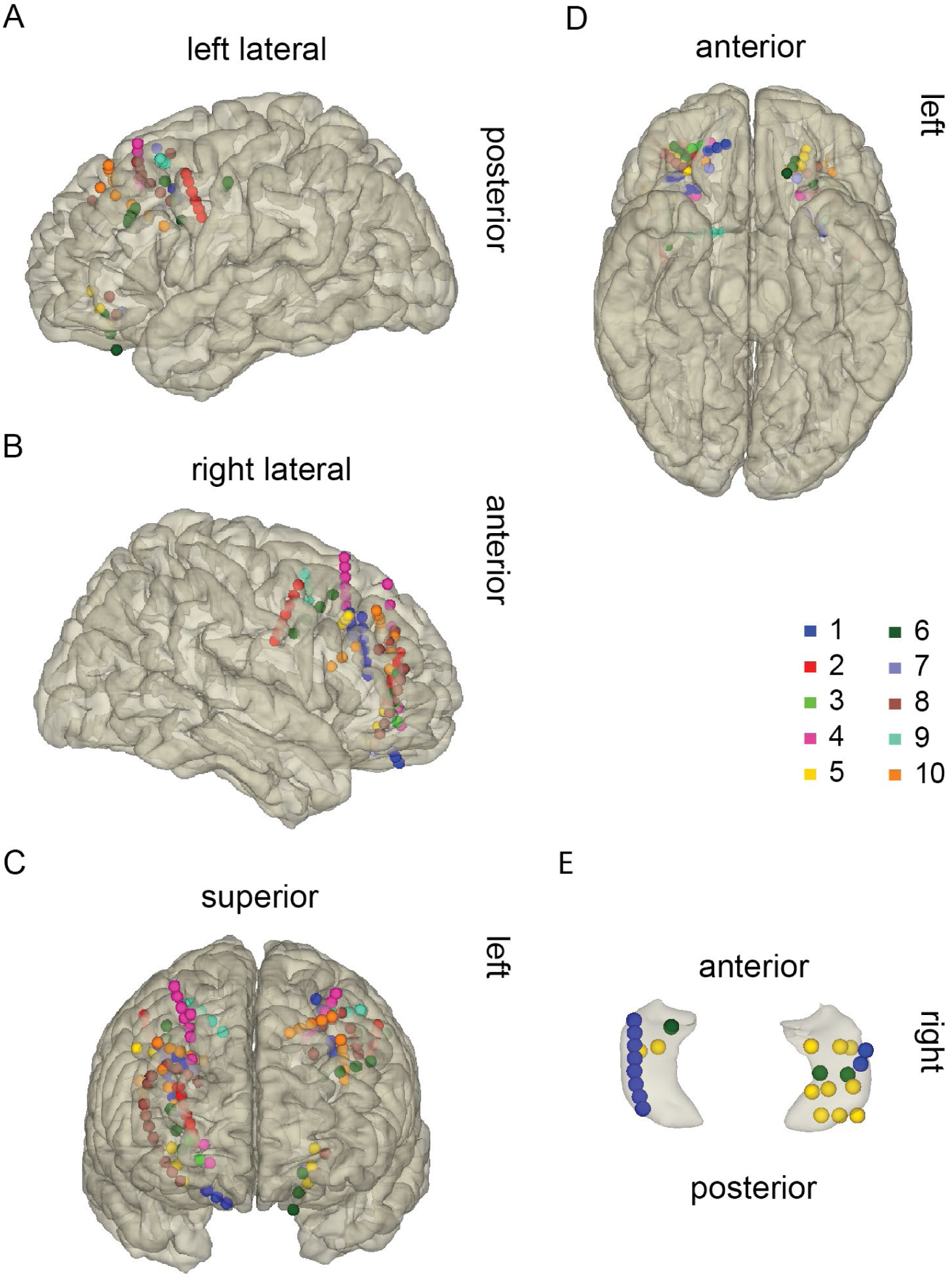
Patients’ electrode coverage is color-coded by the patients’ number. (A-D) shows the PFC and OFC coverage from (A) the right sagittal, (B) the left sagittal, (C) the coronal, and (D) the inferior view. (E) shows the hippocampal coverage in a 3D glass hippocampus from the superior anterior view.

#### Behavioral experiment

A control group of 80 healthy adults participated in a movie segmentation test (53 female, mean age = 28, SD = 14, age range = 18 to 68 years old). 31 participants were 18 to 21 years old, 29 participants were 21 to 30 years old, and 20 participants were older than 30 years old.

### Experimental design

Salient events can occur frequently in a flow of events, such as serving customers in a busy café, or they can be infrequent, such as driving along a desert highway. Participants watched a short mute animation (∼3minutes long) that had frequent salient events. The movie was a short version of the animation designed by Ali Derakhshi, named “wildlife” or “Hayat-e Vahsh,” the episode on lions. The movie was selected so that participants have not watched it before, the visual angle was kept similar (Figure 1A), and events with various magnitudes of saliency occurred in a short period. Critically, the storyline of the movie changed so we could test the effect of novel versus anticipated salient events (link to the movie: https://www.youtube.com/watch?v=Q_guH9vA0sk).

The movie had an overarching cliché love triangle story. It starts by showing a few animal couples going back and forth in a park (this part gets repetitive after 2 repetitions). Then, there is a small lion that looks heartbroken. The lion sees a lioness, but there is a bigger lion that also wants to meet the lioness. The two lions fight for the lioness’s attention through a series of matches. After each match, the scored is shown on board (this part is repetitive and predictable). However, the score is not immediately shown after an eating contest. After the bigger lion wins the eating contest, it eats the small lion’s food too. Then the score is shown, and the matches continue. The small lion loses the competition and moves on. The end of the movie shows that the small lion meets a lioness again (this part was repetitive). This movie had periods with an anticipated flow. For example, after each game that the bigger lion wins, the scoreboard is shown.

The iEEG group passively watched the muted movie, but the behavioral control group watched the muted movie and concurrently segmented the movies into episodes. We instructed the behavioral group to press a key “whenever something new happened.” We clarified that “we want to segment this movie into episodes.” After the segmentation task, they performed a target detection task with targets displayed at random intervals. Participants were instructed to press a key as soon as they perceived the target. This part of the experiment measured participants’ response time for normalization the timing of event-boundaries across subjects.

### Behavioral analysis

We recorded the timing of keypresses during segmentation and target detection tasks. Participants’ response time during the target detection task was measured as the difference between the target onsets and the responses. We excluded consecutive keypresses for segmentation if the interval was less than 100ms (double key registration). The averaged response time per participant was subtracted from the timing of keypresses for movie segmentation to normalize the timing of event-boundaries. The number of segmented events was accumulated across participants at 1.5-second epochs. The number of events in the 1.5-second epochs reflected the saliency of events (see also Ben-Yakov and Henson, 2018). Accordingly, an event was more salient if more people reported it as an event-boundary, and an event was considered less salient if fewer people marked them as an event-boundary (See Figure 1B for the range of saliency scale). The autocorrelation analysis of the event saliency was tested using the Ljung–Box test that was implemented in R (R Development Core Team, 2014). There was no significant autocorrelation in the saliency magnitude in 1.5 second epochs (chi-squared = 1.922, df = 1, *P* = 0.1656). Likewise, autocorrelation in the epochs of 1.5-second high-frequency activities was negligible, allowing to use permutation test and apply the event saliency magnitude for statistical analysis as outlined below.

A hierarchical clustering of event relationships (Figure 1) was constructed by applying a binary hierarchical clustering algorithm in R (R Development Core Team, 2014) using distances between events. The distance between two events was measured by adding the saliency magnitude of all the events between the two events. Accordingly, the events that had many highly salient events in between them had larger distance than the events with less salient events between them.

### iEEG data collection and preprocessing

Intracranial EEG data were acquired using the Nihon Kohden recording system, analog-filtered above 0.01 Hz and digitally sampled at 5 KHz or 10 KHz. A photodiode recorded the luminance of a corner of the screen to track the timing of the movie presentation. Two independent neurologists selected the electrodes that showed both epileptic activities and epochs with seizure spread. Only electrodes in non-pathological regions were included in the analysis.

All EEG analyses were run in R, Matlab 2015a, and fieldtrip toolbox (Oostenveld et al., 2010) offline. We applied a 2 Hz stopband Butterworth notch filter at 60 Hz line power noise and harmonics and then down-sampled the data to 1 KHz using resample() Matlab function via fieldtrip. The function applies an antialiasing FIR lowpass filter and compensates for the delay introduced by the filter. All electrodes were re-referenced to a neighboring electrode (i.e., bipolar reference). The continuous signal was then cropped in 1.5-second-long epochs with no overlaps. The epochs were bandpass filtered for high-frequency activity (HFA; 50 to 150 Hz) using padding and a Hamming window. The Hilbert transformation was applied to the filtered data for extracting the power.

### Correlation analysis between the HFA and event saliency

The correlation between event saliency and the HFA in each electrode was calculated using the Spearman correlation. For estimating the p-value, we used a non-parametric statistical permutation test because the non-overlapping 1.5-second epochs of HFA were interchangeable. Note that the magnitude of saliency and HFA epochs were not significantly autocorrelated. The null distribution was made from 1000 iterations of surrogated trial labels. In each iteration, the maximum correlation between the HFA and saliency magnitude was taken across all electrodes in a region of interest (namely, the lateral PFC, OFC, and the hippocampus) for each participant. The proportion of HFA-saliency magnitude correlation-coefficients in the null-distribution that was more than the observed correlation-coefficient yielded the non-parametric corrected p-value for the observed correlation. A p-value < 0.05 was considered significant.

### Effects of anticipating salient events

We tested if the HFA-saliency magnitude correlation changed with anticipating salient events using a linear mixed-effect model. We recalculated the correlations between HFA and saliency magnitudes in each brain region in sliding windows of 15 seconds with overlaps of 7.5 seconds (24 bins). We used a linear mixed-effect model to test the effect of novel (n = 13) or anticipated periods of salient events (n = 11; rep = 0 for novel and 1 for repetitive storylines; Figure 4) from the three regions of interest (the correlation coefficients (R) in each electrode region; the hippocampus, OFC, and dorsolateral PFC). We used the linear mixed-effects model in MATLAB (fitlme) to account for the different number of electrodes in each ROI of a subject and the nested effect of time (formulated as R ∼ rep + (rep | subject : electrodes) + (1|rep: time_bin)). ANOVA was applied for the result of F-tests for the fixed-effect term in the linear mixed-effects model.

### Electrode localization and visualization

Electrode locations were reconstructed and visualized in MATLAB using the Fieldtrip toolbox (Stolk et al., 2018). We manually selected electrodes on the post-implantation CT, which was co-registered to the pre-implantation MRI using SPM (Ashburner and Friston, 1997), to maximize the accuracy of the reconstructions. A neurologist identified the electrodes’ locations. We then normalized each patient’s pre-implantation MRI to the MNI152 template brain using SPM to obtained the electrode positions in MNI space (Ashburner and Friston, 1999). If electrode locations in MNI space did not correspond to electrode locations in native (subject) space after normalization (e.g., an electrode is within hippocampus in native space, but appears outside the hippocampus in MNI space after normalization), then electrode locations were manually adjusted to represent their true locations in native space. Electrode locations for bipolar re-referenced channels were calculated as the mid-point between the two electrodes (Burke et al., 2013, 2014; Long et al., 2014). Representations of the cerebral cortex were generated using FreeSurfer (Dale et al., 1999) and representations of the hippocampus were generated from the Desikan-Killiany atlas (Desikan et al., 2006) using Fieldtrip. Brodmann areas were inferred from Bioimage Suite package (http://bioimagesuite.yale.edu/).

## Results

We identified the magnitude of salient events in the movie by studying a separate group of adults (n = 80). This group indicated when during the movie a new episode started (i.e., perceiving an event-boundary). The metric of event saliency magnitude was the proportion of identified event-boundaries in a short time-window of the movie (about 20 frames or 1.5 seconds; Figure 1B; total movie time was 3 minutes). Epochs of 1.5 sec resulted in saliency magnitude of 0 to 0.6 (1 would be the maximum saliency magnitude when every participant agrees that during the same 1.5 seconds an event-boundary occurred). A distance matrix was constructed from the sum of the saliency of events that occurred between pairs of events and was used for binary hierarchical event clustering (see the methods; Figure 1). We tested the hypothesis that the magnitude of event saliency was tracked in the targeted regions. Ranked (Spearman) correlation and non-parametric permutation tests for statistical results were applied. The interchangeable non-overlapping HFA in 1.5-second epochs allowed using non-parametric permutation testing. Cluster-corrected p-values are reported.

We observed that HFA correlated with the magnitude of event saliency that was captured by the behavior of the independent rating group. The neural effect was clustered in dorsolateral PFC (BA 6, 8, 9, and 10; in 5 out of 7 patients; 7 out of 9 patients with lateral PFC electrodes had dorsolateral PFC coverage). The effect was also detected in the hippocampus (BA 54; in 3 out of 3 patients) and the medial OFC (BA 11; in 3 out of 3 patients with medial OFC coverage; 3 out 7 patients with OFC electrodes had medial PFC coverage; Figures 3). See Figure 3 for the correlation coefficient of all electrodes and Tables 2 for statistical results of electrodes that showed a significant correlation (the R-value of electrodes with p >0.05 is color-coded in Figure 3.)

**Table 2.** Statistical results. The table lists the patient’s number, Spearman correlation coefficient r, cluster-corrected p-value, and the MNI coordinates of the electrode in millimeters and RAS: positive x = right, positive y = anterior, and positive z = superior. The electrodes are sorted according to their localization into the hippocampus, OFC, or PFC, as observed in the MRI scan in native space.

| *Patient* | *r* | *p* | *x* | *y* | *z* |
| --- | --- | --- | --- | --- | --- |
| hippocampus |  |  |  |  |  |
| 1 | 0.257 | 0.021 | -32.42 | -17.61 | -13.84 |
| 1 | 0.251 | 0.025 | -32.85 | -33.69 | -5.65 |
| 5 | 0.273 | 0.006 | 22.62 | -15.30 | -18.68 |
| 5 | 0.217 | 0.044 | 35.49 | -16.88 | -14.58 |
| 5 | 0.230 | 0.031 | 22.39 | -29.53 | -9.728 |
| 6 | 0.246 | 0.011 | 25.13 | -21.11 | -13.12 |
| 6 | 0.283 | 0.002 | 33.78 | -20.60 | -11.82 |
| OFC |  |  |  |  |  |
| 1 | 0.305 | 0.003 | 16.26 | 45.28 | -22.26 |
| 1 | 0.271 | 0.005 | 20.11 | 44.06 | -19.80 |
| 4 | 0.263 | 0.004 | 19.01 | 47.55 | -8.65 |
| 6 | 0.410 | < 0.001 | -14.80 | 34.05 | -27.56 |
| 6 | 0.456 | < 0.001 | -17.17 | 36.55 | -20.63 |
| 6 | 0.241 | 0.009 | -18.54 | 39.42 | -12.64 |
| dorsolateral PFC |  |  |  |  |  |
| 2 | 0.280 | 0.012 | -45.65 | -0.92 | 32.36 |
| 4 | 0.350 | 0.003 | 26.82 | 41.96 | 51.74 |
| 4 | 0.391 | 0.001 | 32.04 | 23.65 | 59.05 |
| 4 | 0.352 | 0.003 | 33.25 | 23.40 | 63.07 |
| 4 | 0.349 | 0.003 | -32.37 | 25.08 | 60.27 |
| 6 | 0.369 | 0.001 | 30.23 | 40.62 | -2.91 |
| 6 | 0.279 | 0.013 | 32.44 | 43.05 | 4.80 |
| 6 | 0.602 | < 0.001 | 37.97 | 1.86 | 30.73 |
| 6 | 0.539 | < 0.001 | 37.01 | 7.27 | 36.46 |
| 6 | 0.409 | < 0.001 | 36.53 | 13.01 | 41.85 |
| 6 | 0.308 | 0.005 | 36.32 | 18.54 | 47.24 |
| 6 | 0.268 | 0.022 | -40.57 | 13.40 | 44.16 |
| 6 | 0.302 | 0.005 | -26.79 | 29.02 | 26.59 |
| 6 | 0.291 | 0.005 | -40.40 | 27.15 | 30.77 |
| 6 | 0.397 | < 0.001 | -37.94 | -12.53 | 42.31 |
| 6 | 0.465 | < 0.001 | -45.67 | -12.97 | 43.13 |
| 6 | 0.336 | 0.001 | -52.64 | -13.45 | 43.82 |
| 9 | 0.231 | 0.013 | 18.2 | 7.735 | 49.02 |
| 9 | 0.247 | 0.008 | -37.13 | 14.50 | 52.60 |
| 9 | 0.215 | 0.025 | -42.09 | 15.50 | 53.60 |
| 10 | 0.245 | 0.024 | 21.50 | 39.55 | 32.72 |
| 10 | 0.258 | 0.013 | -35.98 | 13.74 | 25.19 |

**Figure 3.**
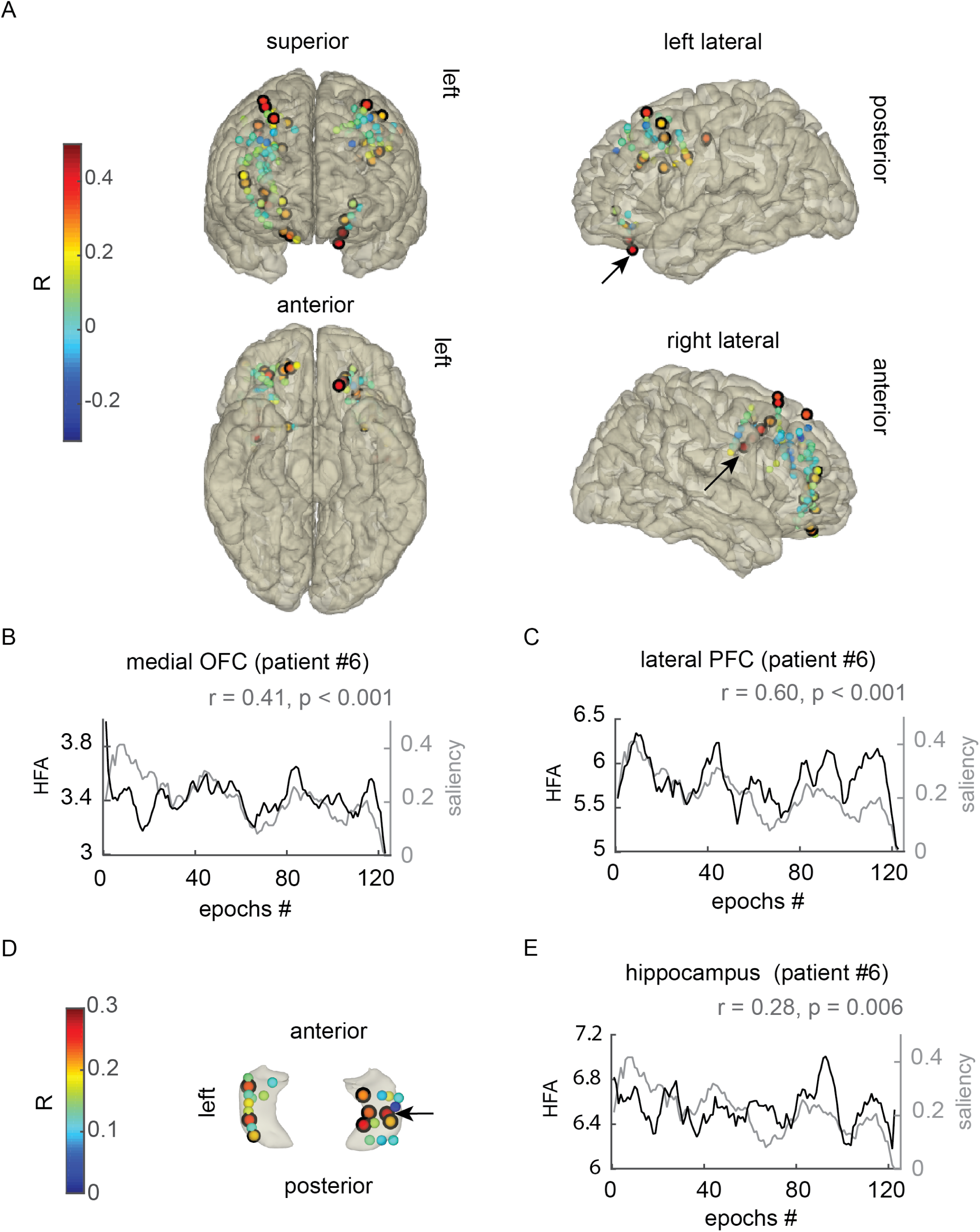
Event saliency was tracked in dorsolateral PFC, medial OFC, and the hippocampus. (A) HFA in the dorsolateral PFC and medial OFC increased with increasing event saliency. The electrodes are color-coded by the Spearman correlation coefficient (R) between the HFA and the event saliency across 1.5-second epochs, which ranged between −0.3 and 0.7. (B) The saliency of each epoch (gray line) and HFA across epochs in a left medial OFC electrode for example (black line). For demonstration, HFA and saliency were smoothed by a 10-episode window (equal to 15 seconds). The underlying data was not autocorrelated. (C) the same as (B) but for in a right lateral PFC electrode. (D) HFA in the hippocampus also correlated with event saliency magnitude. (E) An example of HFA in the hippocampus and the epochs saliency magnitude (same as B and C). (A and D) The black outlines highlight the electrodes that showed the effect (cluster-corrected *P*< 0.05). The thickness of the black outline reflects the effect significance (See Table 2 for the exact p-values < 0.05); the arrows show which electrode corresponds to the plot B, C, and E, which were from patient #6.

We conducted a planned analysis on the electrodes that showed a significant correlation (i.e., task-relevant) to assess the effects of anticipation (Table 2). We re-calculated the correlation coefficient between the HFA and event saliency magnitude in 15 seconds long sliding windows (7.5 seconds overlaps) resulting in 24 tested windows, out of which 11 had repetitive storylines, and 13 were novel. The flow of salient events in 46% of the sliding windows was anticipated. A storyline was predictable if the same type of event reoccurred more than twice, such as repetition of animals going back and forth or repetition of scoring in a competition. The results of a linear mixed effect model showed that the correlation coefficient between HFA and the saliency magnitude was higher in the OFC when the salient events were anticipated than when they were novel (Figure 4A; OFC F(1,142) = 4.3, *P* = 0.039), and this effect was observed in all patients with task-relevant electrodes (Figure 4B). There was no difference between novel and anticipated salient events in the hippocampus (F(1, 166) = 0.39, *P* = 0.52) or the dorsolateral PFC (F(1, 526) = 1.26, *P* = 0.26).

**Figure 4.**
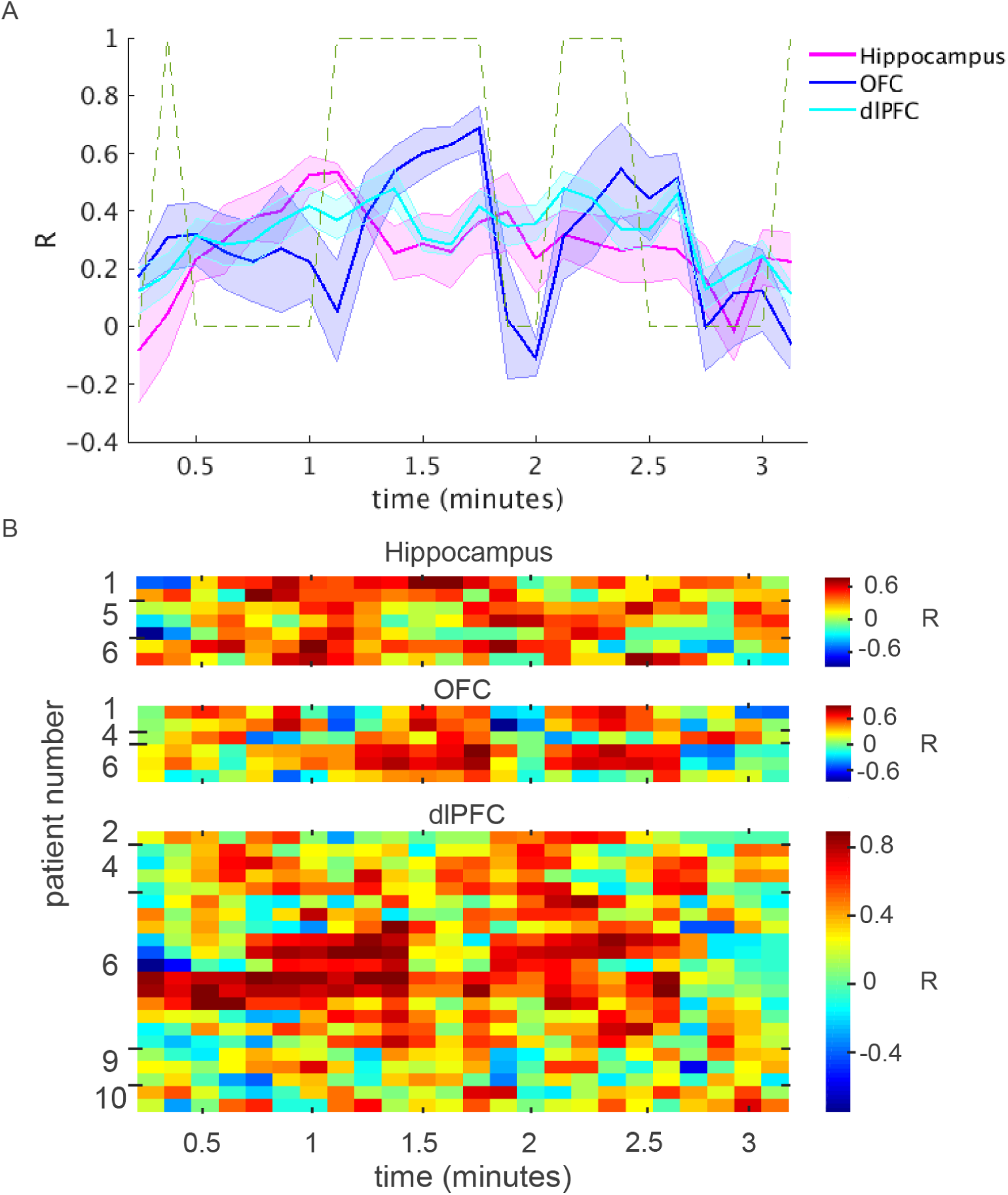
Representation of event saliency during repetitive and novel storylines. The HFA-saliency magnitude correlation coefficients (R) in sliding windows of 15 seconds (with 7.5 seconds overlaps) throughout the movie. (A) The solid lines show the mean R in the hippocampus (in magenta), medial OFC (in blue), and dorsolateral PFC (in cyan), in the y-axis. The x-axis is the time in minutes. The shades lines show the standard error of the mean. The dashed line shows 1 for the periods with repetitive storylines (anticipated salient events) and 0 for novel periods. (B) The R across the time bins is color-coded in each electrode that was included in the planned test. The x-axis is the time in minutes. Each row shows R in an electrode. The patients’ number, showing the owner of the electrode, is written on the y-axis. The R in the OFC was higher for anticipated salient events than for novel events (*P* < 0.05).

## Discussion

During encoding, a sequence of events is segmented to construct a hierarchical representation of event associations (Kurby and Zacks, 2008; Zacks and Swallow, 2007; Zwaan and Radvansky, 1998) with clusters of associated events represented in lower levels of a hierarchy and the associations of the clusters of events represented in higher levels of the hierarchy. The construction of such a hierarchical association requires linking relevant events and separating events that occur in different circumstances. For instance, a circumstance changes with perceiving a deviant event. Detecting the magnitude of event saliency can also contribute to establishing the structure of associations. When the newly perceived event is not salient, the event is closely associated with the preceding events; however, if the new event is highly salient, it should be separated from the preceding events. Here, we report distributed neural regions that detect the magnitude of deviance (i.e., event saliency) in a flow of events including dorsolateral PFC and hippocampus and further show that anticipating the deviant events affects the OFC activities.

We used event segmentations of a large control population (behavior group) who watched the silent movie, to infer the event segmentation in another group with intracranial electrodes (iEEG group) who watched the same movie. The behavior group’s event segmentation provided the event saliency of the entire movie. The reaction time of each subject in this group was estimated from a target detection task and used for normalizing the timing of event-boundaries (see the methods). We inferred the saliency from the proportion of people that reported an event-boundary in each movie epochs. The 1.5 seconds windows provided interchangeable epochs of data for using correlation and permutation tests (see the methods).

The iEEG group passively watched a movie and did not know about the segmentation task, allowing us to study spontaneous and naturalistic neural processing during parsing a continues flow of events. We observed that the HFA, that is linked to nearby SUA activity (Belitski et al., 2008; Jacobs and Kahana, 2009; Lachaux et al., 2012; Ray et al., 2008a; Rich and Wallis, 2017) increased proportionally with event saliency in hippocampus, dorsolateral FPC, and medial orbitofrontal cortex.

HFA in the dorsolateral PFC in the iEEG group tracked event saliency magnitude. Dorsolateral PFC is critical for guiding attention (Corbetta and Shulman, 2002; Hopfinger et al., 2000; Kastner et al., 1999; Paus, 1996) and increased HFA may in part be due to attention to novel events (Ray et al., 2008b; Zacks et al., 2001). Dorsolateral PFC is also engaged in cognitive control and conflict monitoring (MacDonald et al., 2000; Miller and Cohen, 2001) by detecting new associations of categories and exemplars (Dolan and Fletcher, 1997). Accordingly, the observed additional correlation between HFA and event saliency in dorsolateral PFC reflects the demand for event segmentation and updating the event circumstance (Reynolds et al., 2007; Zacks et al., 2007).

Event saliency also correlated with HFA in the hippocampus. Hippocampal activity has been linked to the representation of event associations (Ekstrom et al., 2003; Mack et al., 2017; Quiroga, 2012; Quiroga et al., 2005). The hippocampal representation changes with salient changes in the environment (Shapiro et al., 1997), and its activity increases with detecting salient events (Axmacher et al., 2010; Chen et al., 2013, 2015; Knight, 1996; Kumaran and Maguire, 2007; Lisman and Otmakhova, 2001; Wittmann et al., 2007). Recent studies showed that hippocampal representations reflect the scale of topological saliencies of an environment, such as changes in the spatial closeness of streets or the centrality of the streets (Javadi et al., 2017), and the scale of deviance from expectation (Chen et al., 2015). Also, the hippocampal BOLD signal tracked the saliency of event-boundaries when people watched movies (Ben-Yakov and Henson, 2018). Here, we propose that small deviance induces only a minor change in the hippocampal representation so that close events share more similar hippocampus representations than far events (Ezzyat and Davachi, 2014). These result expand the pattern separation mechanism for distinguishing similar visual associations attributed to the hippocampus (Yassa and Stark, 2011), to a mechanism for identifying the scale of event separation. Pattern separation for visual stimuli engages dentate gyrus in the hippocampus (Baker et al., 2016; Berron et al., 2016) but what subregion of the human hippocampus contributes to the deviant detection is unknown (see Lisman and Grace, 2005 for the novelty signal in the rodent’s subiculum, and Knierim and Neunuebel, 2016 for mismatch signal in subregions of rodent’s hippocampus).

We also observed a similar saliency magnitude effect in the medial OFC (BA 11 but not in the lateral OFC), with increased HFA for highly salient events. This observation is akin to the representation of saliency in the non-human primates’ OFC, captured by HFA (Rich and Wallis, 2016, 2017). In humans, breaching expectations increases the OFC activity (Duarte et al., 2009; Mikutta et al., 2015; Nobre et al., 1999). OFC also represents the saliency of anticipated events (Bechara et al., 1996; Metereau and Dreher, 2015). The reflection of the saliency magnitude suggests that OFC represents the structure of the event association. Notably, anticipation is a critical feature for encoding sequences of events because the gist of previous experience shapes the expected context (Purcell, 1986; Reynolds et al., 2007). Here, the HFA in medial OFC tracked the event saliency better when the salient events were anticipated than during a novel intrusion of events suggesting that the anticipated structure of event associations is represented in the medial OFC.

An important question concerns the dynamics of interaction in the neural network for representing the structure of event associations. For example, disturbing the input from the hippocampus to OFC impairs representing task structures in rodents (Wikenheiser et al., 2017). It is not clear if hippocampal deviancy detection is essential for constructing the OFC signal in humans or if other brain regions such as mid-brain structures contribute to detecting the magnitude of deviance (Dürschmid et al., 2016; Wittmann et al., 2007). For instance, hippocampal response to deviant events is associated with activity in the substantia nigra and ventral tegmental area (Murty and Adcock, 2014; Wittmann et al., 2007). It is also suggested that the hippocampus provides a novelty signal to the nucleus accumbens (Dürschmid et al., 2016). While hippocampus activity is linked to anticipation (Hsieh et al., 2014; Hindy et al., 2016; Jafarpour et al., 2017), an outstanding question is whether the prediction is made by the hippocampus or is under control of other brain regions such as the PFC. Systematically comparison of regional activity, however, requires simultaneous recordings from both regions in a patient. A caveat of this iEEG study is that all the patients did not have sufficient coverage from multiple task-relevant regions to definitively study the dynamics of the network.

Event segmentation requires tracing the association of a new event to the preceding events (Zwaan and Radvansky, 1998). Detecting a new event’s deviance magnitude helps with an accurate association of the event to the preceding sequence. Accordingly, segmented sequences that are separated by small surprises are more associated in comparison to a sequence separated by a big surprise. Detecting deviant events is known to increase neural activity in the PFC and the hippocampus (Axmacher et al., 2010; Bunzeck et al., 2010; Knight, 1996; Kumaran and Maguire, 2007; Long et al., 2016; Strange et al., 2005). Both brain regions, albeit differently, represent the associations in an experimental setup (O’Keefe and Nadel, 1978; Tolman, 1948; Wikenheiser and Schoenbaum, 2016; Wilson et al., 2014). Here, we showed that the hippocampus and PFC regions tracked the scale of event saliency in a movie, and in the medial OFC this effect is stronger when the salient events were anticipated than for novel events. We propose that a core function of the hippocampus, dorsolateral PFC, and medial orbital PFC network is to construct the event association structure, akin to a task structure (Wikenheiser and Schoenbaum, 2016; Wilson et al., 2014).

## Acknowledgments

The authors are indebted to the patients for their participation. We thank Jie Zheng and members of the Knightlab for helping with data collection. We also thank Prof. Elizabeth Buffalo for helpful discussions. This research used statistical consulting resources provided by the Center for Statistics and the Social Sciences, University of Washington. This work was sponsored by the James S. McDonnell Foundation and NINDS R37 NS21135 to RTK and the UC Irvine School of Medicine Bridge Fund to JJL.

